# Ketamine and sleep modulate neural complexity dynamics in cats

**DOI:** 10.1101/2021.06.25.449513

**Authors:** Claudia Pascovich, Santiago Castro-Zaballa, Pedro A.M. Mediano, Daniel Bor, Andrés Canales-Johnson, Pablo Torterolo, Tristan A. Bekinschtein

## Abstract

There is increasing evidence that level of consciousness can be captured by neural informational complexity: for instance, complexity, as measured by the Lempel Ziv (LZ) compression algorithm, decreases during anesthesia and non-rapid eye movement (NREM) sleep in humans and rats, when compared to LZ in awake and REM sleep. In contrast, LZ is higher in humans under the effect of psychedelics, including subanesthetic doses of ketamine. However, it is both unclear how this result would be modulated by varying ketamine doses, and whether it would extend to other species. Here we studied LZ with and without auditory stimulation during wakefulness and different sleep stages in 5 cats implanted with intracranial electrodes, as well as under subanesthetic doses of ketamine (5, 10, and 15 mg/kg i.m.). In line with previous results, LZ was lowest in NREM sleep, but similar in REM and wakefulness. Furthermore, we found an inverted U-shaped curve following different levels of ketamine doses in a subset of electrodes, primarily in prefrontal cortex. However, it is worth noting that the variability in the ketamine dose-response curve across cats and cortices was larger than that in the sleep-stage data, highlighting the differential local dynamics created by two different ways of modulating conscious state. These results replicate previous findings, both in humans and other species, demonstrating that neural complexity is highly sensitive to capture state changes between wake and sleep stages while adding a local cortical description. Finally, this study describes the differential effects of ketamine doses, replicating a rise in complexity for low doses, and further fall as doses approach anesthetic levels in a differential manner depending on the cortex.

**In brief:** Previous studies have shown that Lempel Ziv complexity (LZ) decreases during anesthesia and non-rapid eyes movement (NREM) sleep in humans and rats whereas it increases in REM sleep and under the effect of psychedelics. In this work we show that in the cat, LZ is lowest in NREM sleep, but similar in REM and wakefulness. We also found a ketamine inverted U-shape dose-response curve only in the auditory and prefrontal cortex, with a much larger variability in the ketamine across cats and cortices when compared to the sleep cycle.

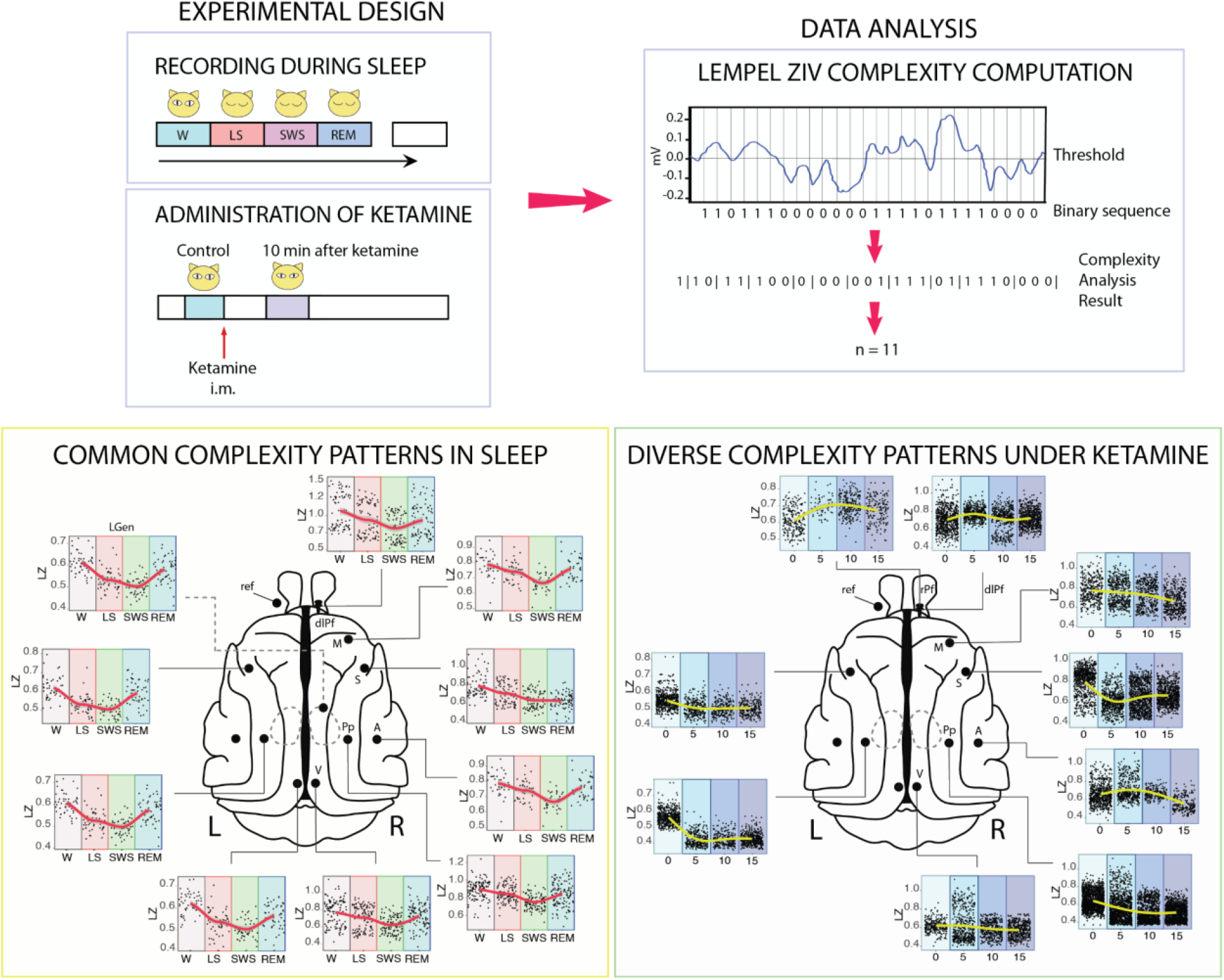

## Introduction

There is increasing evidence for a strong association between neural information measures, such as electrophysiological signal complexity, and level of consciousness (Abásolo et al., 2015; Castro-Zaballa et al., 2019; Mateos et al., 2018; Schartner et al., 2015; Schartner, 2017; Zhang et al., 2001). One of the most studied neural complexity metrics is Lempel-Ziv complexity (LZ), capturing the number of distinct substrings or patterns within a sequence (Lempel & Ziv, 1976; Ziv & Lempel, 1978). A decrease in complexity has been demonstrated for anesthesia (Li & Mashour, 2019; Schartner et al., 2015; Zhang et al., 2001), and during non-rapid eye movement sleep (NREM sleep) when compared to normal wakefulness. However, REM complexity has consistently been shown to be above NREM sleep and below normal wakefulness (Abásolo et al., 2015; Andrillon et al., 2016; Mateos et al., 2018; Schartner et al., 2017). The increase in complexity during REM, where vivid dreaming often occurs, may lend credence to the hypothesis that complexity may not only be modulated by consciousness level but also signal the degree of contents of consciousness (Abásolo et al., 2015; Mateos et al., 2018).

Further evidence for LZ associated with an increase in the range of conscious contents comes from higher LZ during resting state in humans under the effect of psychedelics, specifically lysergic acid diethylamide (LSD), psilocybin, and subanesthetic doses of the dissociative NMDA-antagonist ketamine, compared to placebo (Li & Mashour, 2019; Mediano et al., 2020; Schartner, et al., 2017). These drugs have profound and widespread effects on conscious experiences, both internally and externally generated. More specifically, they appear to “broaden” the scope of conscious contents, vivifying imagination and positively modulating the flexibility of cognition (Carhart-Harris et al., 2016; Carhart-Harris et al., 2014). For all three drugs, reliably higher spontaneous signal diversity was reported. More recently, a higher level of complexity following a subanesthetic dose of ketamine was also reported (Farnes et al., 2020; Li & Mashour, 2019) in spontaneous high-density scalp electroencephalography (EEG) signals in healthy volunteers, but no increase was observed when auditorily stimulated.

Ketamine also appears to maintain spatiotemporal complexity, as measured through the perturbational complexity index (PCI) (Sarasso et al., 2015). PCI is the result of applying LZ to the spatiotemporal pattern of cortical activation evoked by transcranial magnetic stimulation (TMS), and has proven to be a reliable classifier of level of consciousness (Casali et al., 2013). PCI decreases during propofol, midazolam and xenon anesthesia (Casali et al., 2013), but maintains wakefulness baseline level during ketamine anesthesia (Sarasso et al., 2015).

Despite this body of work, important questions remain unanswered. First, prior studies provide only a disjointed picture by investigating the effect of anesthetic dose in TMS-evoked cortical activation (Sarasso et al., 2015) or subanesthetic dose in spontaneous magnetoencephalographic (MEG) signals (Schartner et al., 2017). For a more complete understanding of ketamine’s psychoactive effects, a systematic investigation of the dose-dependent effects of ketamine on cortical complexity using the same modality is required. Therefore, in this work we aimed to investigate the level of informational complexity during different stages of sleep in the cat as well as under subanesthetic doses of ketamine in a dose-dependent manner, compared to the control awake state. Additionally, we determined how the complexity measures under ketamine compared to baseline conditions, with or without the presence of sensory stimulation. Finally, we sought to understand the possible differences in informational complexity between resting-state periods and sensory stimulation periods across conscious states. We aim to add to the characterization of the interaction between psychedelic states and perturbational states in intracranial recordings and via dose dependent manner since our own work suggests a modulation by task (Mediano et al. 2020; Mediano et al, 2021) while others don’t (Farnes et al., 2020). Accordingly (Pascovich et al., 2019), the following hypotheses were proposed: (1) LZ would reflect sleep level: LZ in wakefulness would be just above REM sleep. REM sleep would be above light sleep (LS), and NREM sleep would have the lowest complexity value; (2) LZ would be increased during the initial period of drug infusion compared to baseline wakefulness; (3) the level of complexity would be higher under sensory stimulation compared to baseline, for both conditions, with and without ketamine; and (4) stimulation-induced complexity increase would be more evident under the effect of ketamine. This last hypothesis is line with the entropic brain theory, assuming that under psychedelics the diversity of mental states is increased and the experience produced by a stimuli is amplified by the brain under this state (Carhart-Harris et al., 2016; Carhart-Harris et al., 2014).

## Results

### Sleep shows a state-dependent effect on Lempel-Ziv complexity

Cats underwent a polysomnographic recording in semi-restricted conditions where they were adapted to sleep. Data were obtained during spontaneously occurring quiet wakefulness, LS, NREM sleep and REM sleep (Figure 1A). Examples of raw traces and power spectrum characterization in two cortices from one cat are presented in Figure 1D. For a more detailed description of power spectral density results see Castro et al. (2019).

**FIGURE 1.**
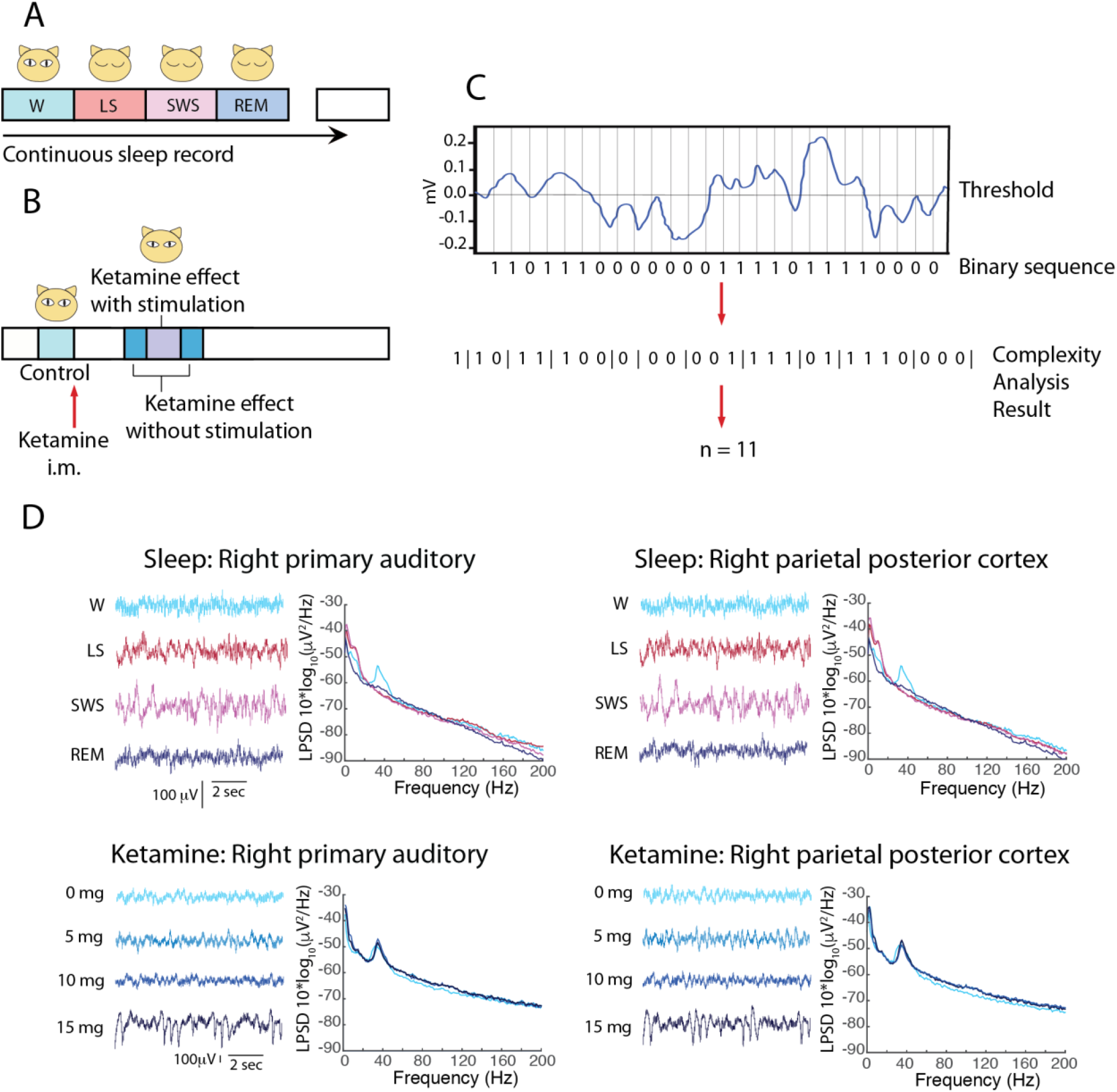
Schematic illustrating the experimental design for electrocorticographic recordings during the different states of sleep (A) and before, and after the different doses of ketamine (B). i.m., intramuscular. (C) Illustration showing how to transform a segment of ECoG signal series into a binary sequence and the result of the LZ complexity analysis on the binary sequence. (D) Raw traces and spectral characterization of sleep stages and different subanesthetic doses of ketamine in one animal.

LZ was computed using the LZ78 algorithm (Ziv & Lempel 1978; Figure 1C) from the different sleep stages for all the cortices available (Figure 2A). Effect sizes for differences between states at the single subject level are shown in Figure 2B. For all animals, LZ scored higher for wakefulness than NREM sleep (Cohen’s d > 0.8) for most of the cortices. As predicted, LZ values were highest for REM and W, intermediate for LS, and lowest for NREM.

**FIGURE 2.**
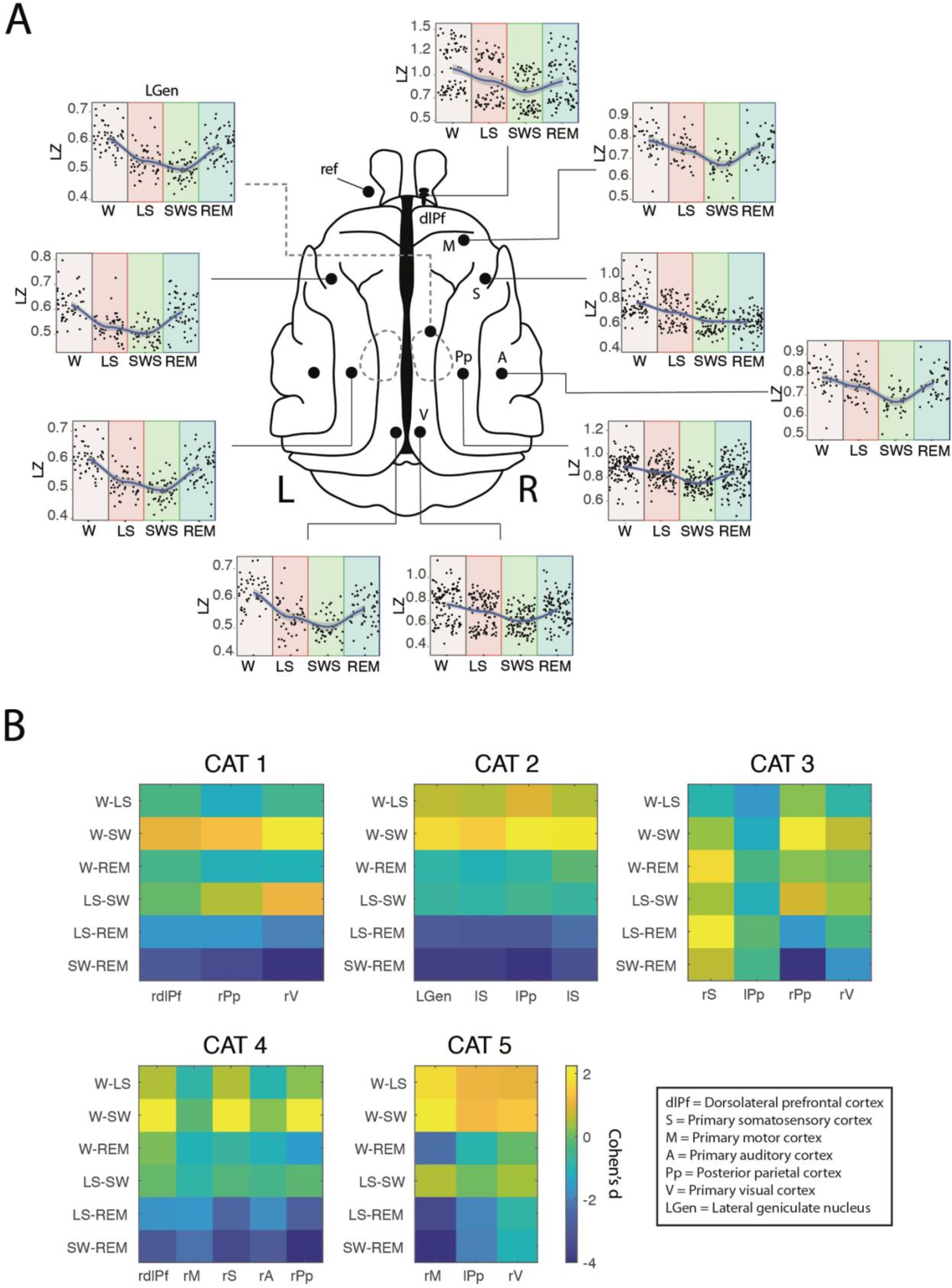
Cortical dynamic of LZ during sleep. Schematic representations of the cat brain are used to visualize the differential dynamics of LZ during wakefulness and different states of sleep (A), showing a U-shaped complexity curve with state progression from W to LS, SWS and REM. “L” indicates Left side and “R” right side. (B) The differences in average LZ between sleep states, as measured by ANOVA and Tukey post-hoc test. Effect sizes were calculated by Cohen’s d and represented in a colour scale, where yellow means a positive difference and blue means a negative difference between the effect sizes of the pair of states compared. W = wakefulness; LS = light sleep; SWS = slow wave sleep; REM = rapid eye movements sleep; dlPf = dorsolateral prefrontal cortex; Pp = posterior parietal cortex; V = visual cortex; LGen = lateral geniculate nucleus; S = somatosensory cortex; M = motor cortex; A = auditory cortex; ref, reference electrode location. In Figure B, “r” indicates right and “l” indicates left cortex.

Additionally, mixed effects models were formulated for each cortex including the cat as a random effect when applicable. Thereafter, model selection was performed between linear and quadratic models using Bayes Factors (BF) to decide between U-shaped and linear fits. All model comparisons between linear and non-linear quadratic fits showed the supremacy of the non-linear fit (Table 1) in agreement with our previous hypotheses, where the REM sleep showed higher complexity values than the deep sleep - with the exception of the right somatosensory cortex, where the results showed a flattening of the curve compared to all other cortices (Figure 2).

**TABLE 1.** Selection between linear and non-linear models among different sleep stages for each cortex. Mixed effects models were formulated for each cortex including the cat as a random effect when applicable. Bayes Factors (BF) were used to decide between U-shaped and linear fits. With the exception of the right primary somatosensory cortex, all model comparisons showed the supremacy of the quadratic fit. The asterisks indicate substantial evidence for a quadratic fit (BF >5).

| Cortex | Nº of cats | Model | BF |
| --- | --- | --- | --- |
| Right dorsolateral prefrontal | 2 | linear |  |
|  |  | quadratic * | 6.88x10<sup>18</sup> |
| Right primary motor | 2 | linear |  |
|  |  | quadratic * | 1.90x10<sup>17</sup> |
| Right primary auditory | 1 | linear |  |
|  |  | quadratic * | 4.42x10<sup>6</sup> |
| Right primary somatosensory | 2 | linear |  |
|  |  | quadratic | 0.681 |
| Left primary somatosensory | 1 | linear |  |
|  |  | quadratic * | 8.53x10<sup>20</sup> |
| Right posterior parietal | 3 | linear |  |
|  |  | quadratic * | 1.76x10<sup>7</sup> |
| Left posterior parietal | 1 | linear |  |
|  |  | quadratic * | 7.19x10<sup>9</sup> |
| Right primary visual | 3 | linear |  |
|  |  | quadratic * | 3.70x10<sup>11</sup> |
| Left primary visual | 1 | linear |  |
|  |  | quadratic * | 3.77x10<sup>17</sup> |
| Right Lateral Geniculate | 1 | linear |  |
|  |  | quadratic * | 7.00x10<sup>20</sup> |

### Heterogeneous cortical dynamics across cortices under ketamine

For this experiment, the data were collected under the same experimental conditions as for sleep recordings in the same cats, and i.m. injections of ketamine of 5, 10 or 15 mg/Kg were performed in separate non-consecutive days as schematized in Figure 1B (see Methods). The raw data shown in Figure 1D reveals the presence of slow waves with 15 mg/Kg dose of ketamine, whereas the power spectral density plots show an increase in gamma power, as already had been reported by Castro et al. (2019). Again, LZ was calculated in epochs before and after the administration of the drug. To address dose-response relationships, a multilevel model was used where LZ was predicted by dose (fixed effect), and cat and session were considered as random effects (with sessions nested within cats, and each dose of ketamine was repeated four times).

Considerably greater LZ variability was observed under ketamine than for the sleep results, especially during the lowest doses explored (Figure 3A). In some regions, the results are in agreement with our hypothesis, which predicted an increase in informational complexity after the lowest ketamine dose, followed by a decrease with the higher dose showing an inverted U-shaped relationship. This can be observed clearly in the right rostral and dorsolateral prefrontal cortices as well as the right primary auditory cortex (BF = 1.70×10^14^, BF = 2.57×10^5^, BF = 8.25×10^9^, respectively). However, when we look at the individual effect per cortex in each animal, it can be seen that the inverted U-shaped relationship is not systematic between cats and is present only in the cortices of 2 out of 5 cats (Figure 3B).

**FIGURE 3.**
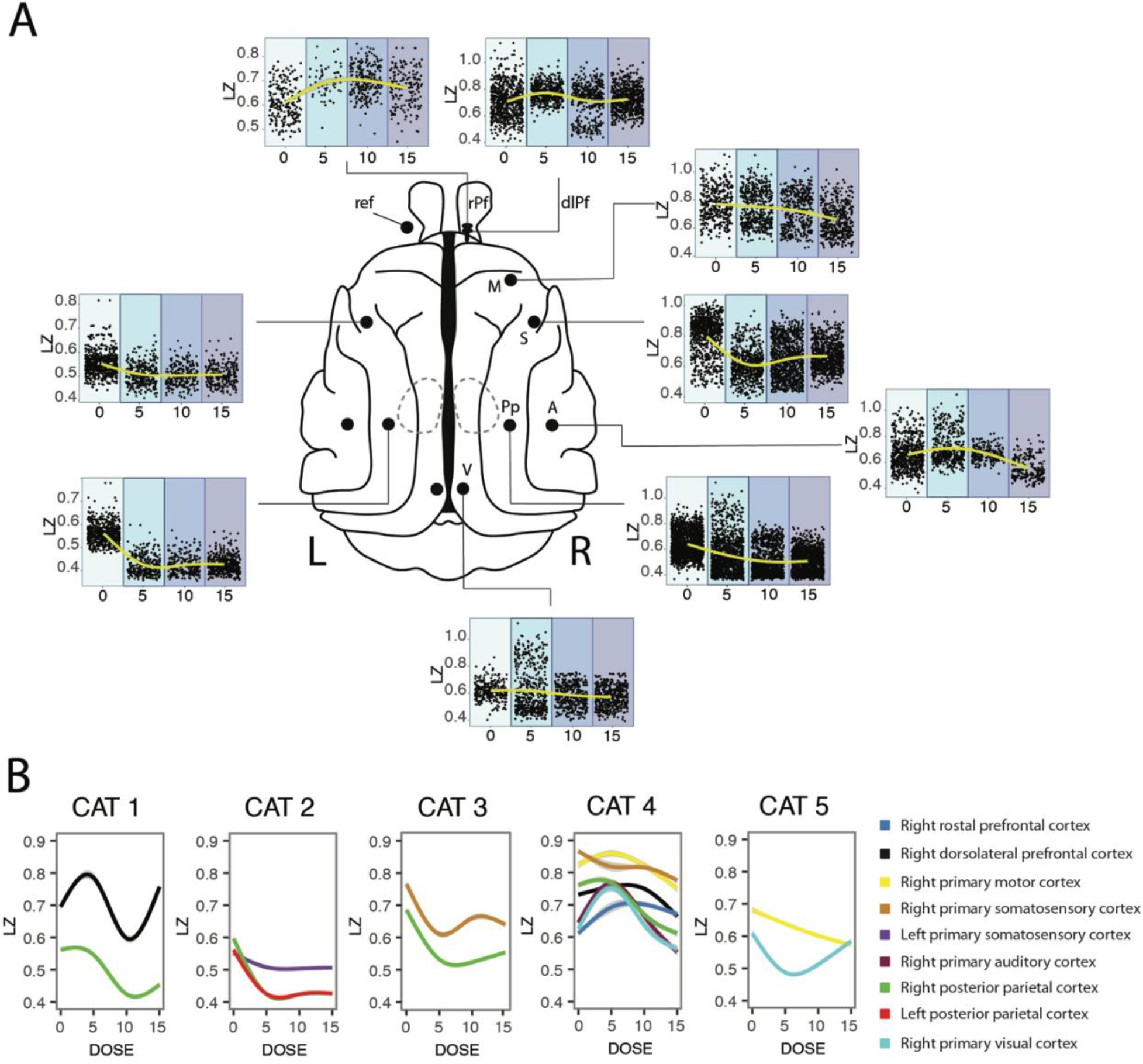
Curves dose-response of the dose of ketamine on cortical dynamics of LZ. (A) Dose-response curve of subanesthetic doses of ketamine, showing an inverted U-shaped curve only for prefrontal and auditory cortices, with monotonic decrease of complexity with concentration for the other cortices. Each plot represents the sum of the different sessions for each dose of the different cats which have that cortex, therefore the Nº of cats is different per cortex. (B) The curves are plotted per cat. It can be clearly seen that the variability in the informational complexity dynamic per cortex and per cat is evidenced more clearly when plotted individually, and shows that there is a dissociation of the dose-response and the anatomical location. The doses are represented in mg/Kg. rPf, rostral prefrontal cortex; dlPf, dorsolateral prefrontal cortex; M, primary motor cortex; S, primary somatosensory cortex; A, primary auditory cortex, Pp, posterior parietal cortex; V, visual cortex. “L” indicates the left side and “R” the right side.

On the other hand, an opposite curve was obtained for somatosensory and posterior parietal cortices. Finally, for the visual cortex the effects were less consistent among cats; in this last example, the two cats tested had different responses to ketamine with opposite effects (Figure 3B). As for sleep, we studied ketamine effects on LZ using model fitting of the individual mixed effects models for each cortex. Model selection was performed in this case between linear, quadratic and cubic models using BF (Table 2).

**TABLE 2.** Selection between linear and non-linear models among different doses of ketamine for each cortex. Mixed effects models were formulated for each cortex including the cat as a random effect when applicable. Bayes Factors (BF) were used to decide between quadratic (U-shaped), cubic and linear fits. Clear evidence towards a quadratic fit was found for right dorsolateral and rostral prefrontal cortices, right primary auditory cortex and left posterior parietal cortex. The asterisks indicate substantial evidence for a quadratic fit (BF >5).

| Cortex | N° of cats | Model | BF |
| --- | --- | --- | --- |
| Right dorsolateral prefrontal | 2 | linear |  |
|  |  | quadratic * | 2.57x10<sup>5</sup> |
|  |  | cubic | 288.73 |
| Right rostral prefrontal | 1 | linear |  |
|  |  | quadratic * | 1.70x10<sup>14</sup> |
| Right primary motor | 2 | linear |  |
|  |  | quadratic | 0.04 |
| Right primary auditory | 2 | linear |  |
|  |  | quadratic * | 8.25x10<sup>9</sup> |
| Right primary somatosensory | 3 | linear |  |
|  |  | quadratic | 3.0x10<sup>-3</sup> |
|  |  | cubic | 0.66 |
| Left primary somatosensory | 1 | linear |  |
|  |  | quadratic | 1.08 |
| Right posterior parietal | 4 | linear |  |
|  |  | quadratic | 1.13x10<sup>-4</sup> |
| Left posterior parietal | 2 | linear |  |
|  |  | quadratic * | 46057.12 |
| Right primary visual | 2 | linear |  |
|  |  | quadratic | 1.28 |
|  |  | cubic | 3.0x10<sup>-3</sup> |

Finally, in order to show the possible inter-areal differential effects of ketamine and dependencies to LZ in basal conditions among cortices, we studied both the baseline variance and its change per area. To study the basal conditions among cortices we built a linear mixed effect model including cats and sessions as random effects, and cortex as fixed effect and that model is statistically reliable (BIC = −15175) when contrasted against a null model (BIC = −12941; p < 0.01) indicating that LZ vary among cortices. We further show that the effect of ketamine does not seem to be dependent on the LZ in basal conditions in wakefulness (see Supplementary Figure 1). When looking into wakefulness and light sleep the effects in LZ were all in the same direction and comparable in intensity independently of the baselines variability. It seems that basal LZ does not predict whether the LZ increases or decreases with drug. Finally, an interesting exploratory finding showed higher baseline LZ for right parietal cortices when compared to left side ones (p < 0.01; BF = 3.3×10^16^), but not strong enough for primary somatosensory and primary visual.

### Informational complexity is not modulated by auditory stimuli under ketamine

In 3 cats, modulation by auditory stimuli was studied. Under control conditions without ketamine, an increase in LZ was observed during stimulation in dorsolateral prefrontal (0.66±0.04 to 0.70±0.007, p < 0.01, η^2^ = 0.044, BF = 10619.07) and auditory (0.63±0.05 to 0.66±0.007, p < 0.01, η^2^ = 0.035, BF = 75.27) cortices, whereas the effect on other cortices studied were non-reliable, including right posterior parietal cortex (0.53±0.02 to 0.53±0.001, p < 0.01, η^2^ = 0.008, BF = 7.0 x10^−5^), right somatosensory cortex (0.62±0.05 vs 0.62±0.002 with p = 0.61, η^2^ = 0.005, BF = 1.0 ×10^−4^), left somatosensory cortex (0.52±0.005 vs 0.53±0.002 with p < 0.01, η^2^ = 0.002, BF = 0.09), and left posterior parietal cortex (0.47±0.01 vs 0.48±0.001, p = 0.85, η^2^ = 0.001, BF = 1.0×10^−4^, Figure 4A).

**FIGURE 4.**
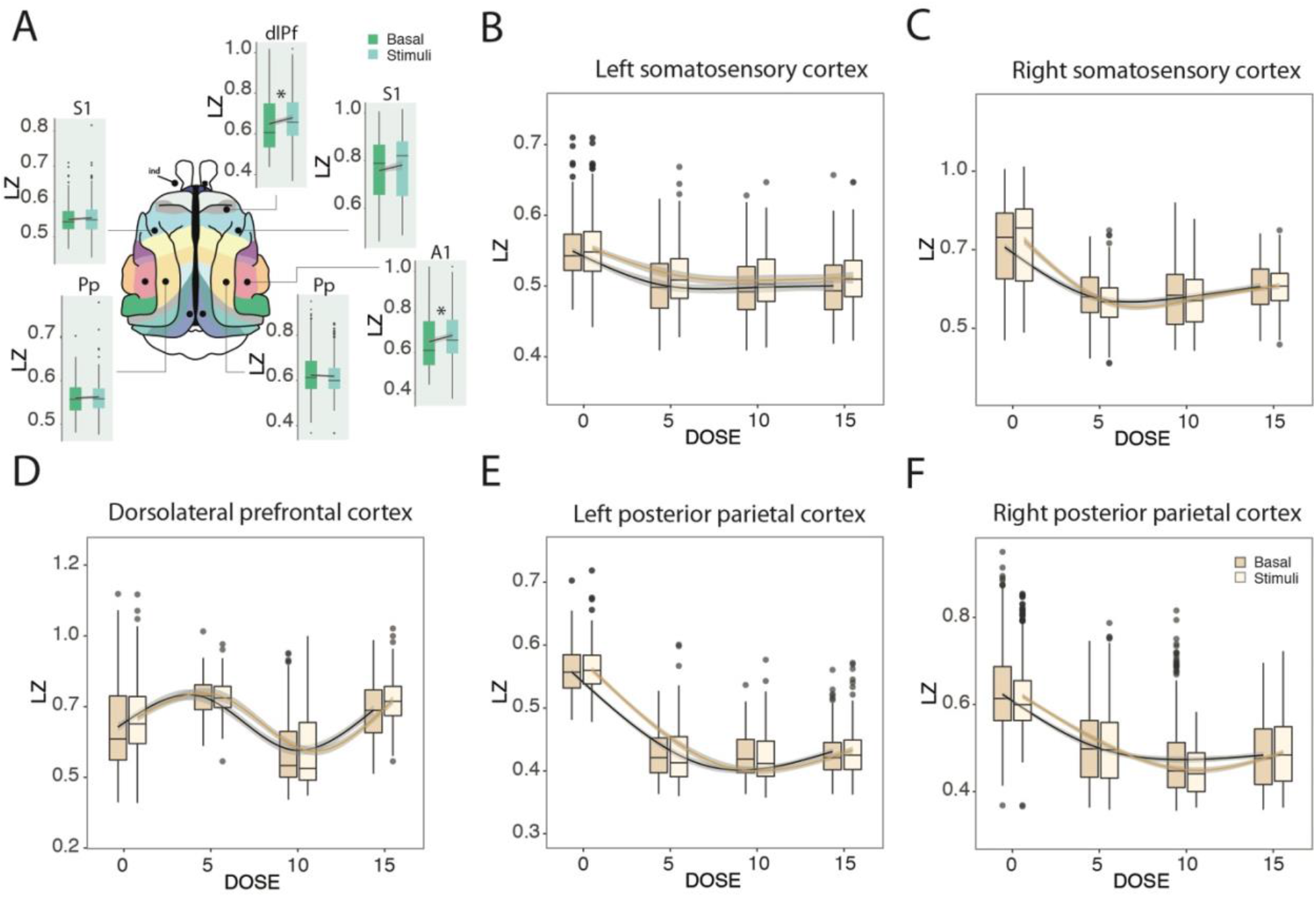
Modulation of LZ by auditory stimulation. (A) The effect of stimulation was shown without ketamine where an increase in LZ was observed during stimulation in dorsolateral prefrontal and auditory cortices. * Statistically reliable (p<0.01; BF > 5). dlPf = dorsolateral prefrontal cortex; Pp = posterior parietal cortex; S1= primary somatosensory cortex; A1= primary auditory cortex. (B-F) Modulation by stimulation under the effect of ketamine in 5, 10 and 15 mg/Kg doses. No stimulation by dose interaction was observed. The doses are represented in mg/Kg.

Initially we hypothesized that the increment in complexity under the sensory stimulation versus non-stimulation conditions would be more evident under the effect of ketamine. However, there was no interaction between stimulation and ketamine. For left somatosensory cortex, non-reliable effect was observed during basal conditions in response to the stimuli (p = 3×10^−4^, BF = 0.09), as well as evidence for no interaction between dose and stimuli (p = 0.16; BF = 3.5×10^−4^, Figure 4B). For right somatosensory cortex, where non-reliable increase was evidenced in control conditions (Figure 4A), no reliable interaction was found during ketamine effect (p = 2.0×10^−3^; BF = 0.012) with no response to the stimuli (p = 0.64; BF = 1.5×10^−4^; Figure 4C). For the prefrontal cortex, where an increase was observed in control conditions, the same effect was found under ketamine (p = 9.4×10^−8^; BF = 335.92), with non-reliable interaction between stimuli and ketamine (p = 0.51; BF = 2.5×10^−8^; Fig. 4D). For left posterior parietal cortex, non-reliable effect was found under baseline conditions, there was no effect of stimulation under ketamine (p = 0.85; BF = 1.2 x10^−4^), and the interaction also remained unchanged under ketamine (p= 0.74; BF = 2.5×10^−6^; Figure 4E). Finally, for right posterior parietal cortex, no effect was found under basal conditions, there was no change with the stimuli under ketamine (p= 0.46; BF= 7.4×10^−5^), and the modulation by stimulation was non-reliable (p = 8×10^−3^; BF = 8×10^−3^; Figure 4F).

## Discussion

In this work, using LZ as a measure of dynamical complexity on direct intracranial recordings, we studied the effect of subanesthetic doses of ketamine in a dose-dependent manner. Ketamine elicited a diverse set of dynamics, with the lower doses showing the most variable effects. For prefrontal and auditory cortices an increase in LZ was observed from low to medium ketamine dose. However, a decrease was evidenced at the maximum dose, drawing an inverted U-shape dose-effect curve, whereas the opposite effect was observed for other cortices including somatosensory and posterior parietal cortices, where an initial decrease was followed by an increase in complexity at higher doses. Additionally, we also presented auditory stimulation to the cats, which elicited an increase in LZ in prefrontal and auditory cortices, but this effect was not modulated by ketamine. Finally, in the same animals, we studied LZ during sleep, which by contrast show an homogeneous pattern among cortices. We demonstrate that informational complexity in the cortex of the cat decreases in light and deep sleep compared to awake states and REM. The same effect was observed in the geniculate nucleus of the thalamus, but as it was tested in only one animal and hence more evidence is required to see convergence in subcortical structures. For most of the cortex, there is only marginal complexity difference between wakefulness and REM sleep. The results were consistent among cats and similar for all the cortices studied and, more importantly, confirm previous results in humans and rats.

As measures of neural signal diversity are known to be sensitive to conscious level in natural state changes (the sleep-wake cycle), they are also sensitive to the changes in brain dynamics associated with psychedelic and anesthetic states. Specifically, Schartner et al. found increased global neural signal diversity for the psychedelic state induced by ketamine, psilocybin and LSD, as compared to placebo, across a range of measures (Schartner et al., 2017). Other recent MEG and EEG studies have also demonstrated elevated signal diversity induced by canonical serotonergic psychedelics and ketamine (Tagliazucchi et al., 2014; Timmermann et al., 2019).

From the perspective of its effects on EEG signal diversity, the dissociative NMDA-antagonist ketamine diverges from traditional anesthetics at subanesthetic concentrations, as it induces dissociative states characterized by a maintained or enhanced repertoire of brain states (Li & Mashour, 2019; Schartner et al., 2017). This is in contrast to GABAergic anesthetics such as propofol, which have been shown to degrade sensory integration and attenuate neural signal diversity in a dose-dependent manner (Ferenets et al., 2006, 2007; Ishizawa et al., 2016). While those studies were based on EEG signals that had been low-pass filtered at 55 Hz and lacked cortical dynamics in higher gamma frequencies, Pal et al. (2020) have recently demonstrated that this part of the signal is important. Using intracranial EEG data from frontal and parietal cortices of rats receiving ketamine or propofol anesthesia, they demonstrated a reduction in broadband (0.5–175 Hz) EEG complexity during ketamine anesthesia that is comparable to that induced by the GABAergic anesthetic propofol. Bandwidth-specific analyses restricted to higher gamma frequencies showed that ketamine anesthesia is distinguished from propofol by suppression of EEG complexity in high gamma frequencies in the range of 65–175 Hz, which previous human studies using scalp EEG could not reveal (Pal et al., 2020). In the present study, by using intracranial electrodes in cats, we were able to study broadband (>0.5Hz) signal complexity.

Contrary to the apparent convergence of psychedelics (LSD, N,N-Dimethyltryptamine or DMT, psilocybin) reported (Schartner et al., 2017), some of us (González et al., 2021) have shown that the effects of ibogaine, a psychedelic alkaloid, induces high gamma power but are less coherent and less complex than control condition, and similar to natural REM sleep. Although some differences in the complexity measure or animal model may explain the difference, it is key to highlight that the ibogaine local complexity patterns were more consistent than those found in the current study, pointing to a different mode of action between alkaloid, serotoninergic and N-methyl-D-aspartate (NMDA) psychedelics.

Ketamine’s primary mechanism of action is as an NMDA antagonist whose receptors are located quite ubiquitously across the cerebral cortex, as well as subcortically (Conti et al., 1994; Huntley et al., 1994). A differential interaction with various subtypes of NMDA receptors could explain the heterogeneity in cortical response under the effects of ketamine (Zanos et al., 2018). However, the non-NMDA receptor effects of ketamine cannot be discounted, in particular its interactions with opioid receptors and hyperpolarization-activated cyclic nucleotide-gated (HCN) channels (Chen et al., 2009; Zanos et al., 2018; Zhou et al., 2013). Additionally, ketamine may indirectly exert effects through its interaction with other circuits. Previous work reported that subanesthetic doses of ketamine increased the release of not only 5-hydroxytryptamine (5-HT) (Amargós-Bosch et al., 2006; López-Gil et al., 2012, 2019), but also noradrenaline (Lorrain et al., 2003) as well as glutamate (Moghaddam et al., 1997) in the medial prefrontal cortex, which may increase signal complexity. At the receptor level, ketamine blocks excitatory NMDA receptors on fast-spiking cortical interneurons more effectively than those on pyramidal neurons. This results in down-regulation of interneuron activity, and decreased gamma aminobutyric acid (GABA) release at the interneuron-pyramidal neuron synapse (Homayoun & Moghaddam, 2007; Seamans, 2008). This decrease in inhibitory tone (decreased GABA release) results in markedly excited pyramidal neurons. It has been proposed that this may explain why ketamine is associated with increased cerebral glucose utilization and blood flow (Langsjo et al., 2005; Långsjö et al., 2004), and increased EEG gamma oscillations (Blain-Moraes et al., 2014; Castro-Zaballa et al., 2019; Ferrer-Allado et al., 1973; Lee et al., 2013; Schwartz et al., 1974) and may also help us understand the changes observed in the complexity of the signal. However, our results show a decrease of LZ in somatosensory and posterior parietal cortices after the lowest dose of ketamine (Figure 3B). As both of these cortices process somatosensory information, our results may be due to a reduction in the somatosensory information influx, as one of the main effects of subanesthetic doses of ketamine is analgesia (Zanos et al., 2018).

A further possible outcome for subanesthetic doses of ketamine effects that we did not find evidence for is the increased locomotion, as found in rats (Hetzler & Wautlet, 1985). An increase in complexity compared to baseline was found for prefrontal and motor cortices, thus, a connection between this regional increase of LZ complexity and putative increased motor activity could be proposed as a possible explanation, however, as reported in our previous publication using the same dataset (Castro-Zaballa et al., 2019), the cats retained muscular tone but hyperlocomotion was not observed in our experiments, nor in previous studies in cats (Ambros & Duke, 2013; Issabeagloo et al., 2011).

The ongoing discussion about complexity as proxy to study integration in different consciousness states oscillates between perturbational and steady state studies. In a perturbational - complementary-study, Arena et al, (2021) quantified the complexity of electrocorticographic responses to intracranial electrical stimulation in rats, comparing wakefulness to propofol, sevoflurane, and ketamine anesthesia using PCI and PCI state-transition (PCI^ST^) (Comolatti et al., 2019). They found ketamine-induced evoked related potentials (ERPs) mixed features with a brief response followed by an OFF period (albeit long-lasting deterministic activations in half of the animals), and the duration of the resulting phase-locked response was close to that of wakefulness.

The time course of PCI^ST^ revealed similarities to wakefulness, but resulted in an overall reduction of complexity. These results from a perturbational study showed a similar feature to our “state” study in that the ketamine induced effects are cortically variable and not consistent between animals. It is, however, difficult to compare this study directly to our results because we used subanesthetic doses of ketamine (5, 10 and 15 mg/Kg), whereas Arena et al., 2021 used anesthetic doses of ketamine (30 mg/kg), so our hypotheses of higher complexity with low doses cannot be addressed in their study. On the other hand, they explored the effect of perturbational complexity index (PCI) which is an electrophysiological metric for the capacity of cortical circuits to integrate information, whereas we studied the effect of auditory stimulation on the naturally occurring and ongoing cortical complexity and hence the complementarity of the findings should be encouraging for the field (Arena et al., 2021).

Neural diversity, assessed by LZ, is an attractive measure because of simplicity, practical applicability, and consistency with both complexity-based (Tononi et al., 2016; Tononi & Edelman, 1998) and entropy-based (Carhart-Harris, 2018; Carhart-Harris et al., 2014) theories of neural integration and consciousness. The measure is also useful in questions regarding local processing as it is computed at the electrode level, thus was able to demonstrate differential effects in distinct thalamic and cortical brain regions. Indeed, according to the dynamic core hypothesis (Tononi & Edelman, 1998) and subsequent theoretical developments such as Information Integration Theory (Tononi et al., 2016), only certain distributed subsets of the neuronal groups that are activated or deactivated in response to a given task are associated with conscious experience, therefore a large cluster of neuronal groups that together constitute, on a time scale of hundreds of milliseconds, a unified neural process of high complexity can be termed the “dynamic core”. In line with this idea, our results could be interpreted as the prefrontal and auditory cortices, where an increase in LZ was observed under the 5mg dose of ketamine, constituting a part of the “dynamic core”, and somatosensory and posterior parietal cortices playing a different role in neural integration. However our results do not necessarily provide strong evidence for the “dynamic core” over other theoretical interpretations such the entropic brain (Carhart-Harris, 2018) or other complexity and consciousness (Sarasso et al., 2021). Since the predictions from most frameworks are less precise and hardly define the specific pattern of results we present. Furthermore, there are complementary approaches to understand information and complexity dynamics, both using state and perturbational experimental and analyses models and frameworks that illustrate the underdeveloped integration of the theories and experiments in this subdiscipline. Another interpretation of the overall ketamine dose-response results, its variance and dynamics, is that it could reflect a level of connection to the external environment interacting with the pharmacological modulation. We however did not systematically assess this behavioral aspect and hence it is difficult to draw conclusions at that level.

Another useful framework for understanding these results is the neuroscience of arousal, including wakefulness, sleep, circadian rhythms, responsiveness and alertness (Bekinschtein et al., 2009; Brown et al., 2011). Sleep shows a clear change in arousal throughout the day cycle; the intensity of the stimuli needed to wake up a person is maximal in deep sleep and lower in light sleep and REM. This pattern partially mimics the results obtained for informational complexity in this study using electrocorticogram recordings (ECoG), and several other nonlinear measures such as fractal dimension and other entropy methods (Ma et al., 2018), but not to other measures such as power in different bands and connectivity methods. This finding allows us to interpret that LZ may index behaviorally defined wakefulness, or arousability by stimuli (Bonnet et al., 1978). Although ketamine is used as an anesthetic and creates unconsciousness in high doses and hence can be framed in terms of consciousness as wakefulness and arousal, the effects at lower doses require a multidimensional framework, able to accommodate neurological symptoms (dizziness, slurred speech), mood modulations, and psychedelic experiences. In principle, if ketamine had the classic profile of a sedative, responsiveness would monotonically decrease (Brown et al., 2011) and a similar profile would be expected for molecular and neural measures. However, ketamine has an interesting profile as it belongs to a group of hypnotics that show hallucinatory capacities and an hormetic or U-shaped curve (Calabrese & Baldwin, 2001) in EEG and blood flow (Cavazzuti et al., 1987; Tsuda et al., 2007). The hormesis of the dose response allows for the comparison of not only conscious level in the sense of wakefulness but in terms of contents of consciousness in low ketamine and REM sleep. From humans we know that the likelihood of increased richness in mental content during the sleep-wake cycle occurs during REM (Windt & Noreika, 2011) after a decrease in NREM (U-shaped); and we know that the richness of mental content, including hallucinations, peaks early with ketamine before decreasing into sedation and anesthesia (Powers et al., 2015) (an inverted U-shaped curve). In both cases the higher levels of content agree with the higher (or recovering) levels of informational complexity as measured by LZ (Abásolo et al., 2015; Mateos et al., 2018; Schartner et al., 2015; Schartner, Carhart-Harris, et al., 2017; Schartner, Pigorini, et al., 2017). In this study, we compare the consistency of the complexity in the cortex in sleep and the diversity in the ketamine challenge as two putatively very different mechanisms of reaching a higher level of content in consciousness.

Recent findings by Mediano et al. (2020) provide strong quantitative evidence on how environmental conditions have a substantial influence on neural dynamics during a psychedelic experience in humans. This work showed how brain entropy is modulated by stimulus manipulation during a psychedelic experience by studying participants under the effects of LSD or placebo, either with gross state changes (eyes closed vs. open) or different stimuli (no stimulus vs. music vs. video). Results showed that while brain entropy increased with LSD in all the experimental conditions, it exhibited largest changes when subjects have their eyes closed, whereas the entropy enhancing effects of LSD were less marked when participants opened their eyes or perceived external stimuli — such as music or video (Mediano et al., 2020). In the present work, we studied the modulation of auditory stimulation on brain complexity in basal conditions and under increasing doses of ketamine in 3 cats using ECoG recordings with the hypothesis of observing a higher level of complexity under stimulation. However, only a slight increase in LZ was evidenced during stimulation in dorsolateral prefrontal and auditory cortices, whereas a complete lack of or very weak effect were found in the other cortices studied (Figure 4). This weak effect may be explained by the low relevance of the stimulus, as it failed to catch the attention of the animals, compared to extremely salient or meaningful stimuli such as music or video. Further evidence that stimulation studies should exploit more complex stimuli also comes from a recent study were TMS pulses also failed to increase complexity in low doses of ketamine in humans (Farnes et al., 2020). Furthermore, Nilsen and collaborators (Nilsen et al., 2019) were unable to demonstrate an influence of attention in LZ complexity after stimulation while we have reported (Mediano et al., 2020) that LZ is modulated when applying different types of stimulation (music and videos). Additionally, we have recently shown (Mediano et al., 2021) that LZ varies with the level of alertness and also depending on the task, not being restricted to measure the level of consciousness but cognitive and attentional demands.

New experiments using more appropriate stimuli in terms of relevance and salience are needed to better address this hypothesis and further the experimental understanding neural dynamics of information theory, complexity and entropy as the system is modulated pharmacologically.

Our sleep results are consistent with previous results in humans (Andrillon et al., 2016; M. Schartner et al., 2017), as well as in rats (Abasolo et al., 2015). However, a closer read shows some differences: Andrillon et al. (2016) reported a small but reliable decrease in LZ during REM sleep compared to the waking state, possibly due to participants engaged in a task during the waking state, whereas the participants in the Schartner et al. study were simply at rest with eyes closed and not engaged or externally driven by task or stimuli. In our study the animals were also at rest but with eyes open and showed a decrease in LZ during LS and further decrease in SWS, which was similar for all cortices (Figure 2A) in line with previous findings (Andrillon et al., 2016; M. Schartner et al., 2017). However, a greater variability was evident for REM sleep state where in some cortices LZ was equal in level of complexity to wakefulness whereas in others it was similar to LS or to SWS (Figure 2B). The complexity pattern among sleep stages observed in the cortex was also evidenced in the lateral geniculate nucleus (Figure 2A), lending clear convergent evidence to the common effects of informational complexity in the brain beyond the cortex for the sleep wake cycle.

In summary, our data demonstrate that there is a dose-dependent ketamine effect on neural complexity. An increase in complexity compared to baseline was found for some cortices (prefrontal, motor, auditory and visual) only in the lowest doses, while the higher dose frequently showed the lowest informational complexity. However, a decrease in complexity was also seen in somatosensory and posterior parietal cortex in the low doses. The heterogeneity of the ketamine effects between cats and cortices contrasts with the homogeneity of the changes in complexity seen for different stages of sleep, further highlighting the differences between natural and pharmacologically induced changes in consciousness. The individual and cortical variability in the neural complexity dynamics revealed by ketamine highlights the intricacy of the brain when altered by dissociatives and psychedelics, pushing for a multidimensional framework beyond simple arousal and alertness parameters to characterize the change in the states of consciousness from a neuropharmacological perspective.

## Methods

### Animals

Five adult cats were used in this study; all of whom were also utilized in a previous report (Castro-Zaballa et al., 2019). The animals were obtained from and determined to be in good health by the Institutional Animal Care Facility of the Faculty of Medicine (University of the Republic, Uruguay). All experimental procedures were conducted in accordance with the Guide for the Care and Use of Laboratory Animals (8th edition, National Academy Press, Washington DC, 2011) and were approved by Institutional and National Animal Care Commissions of the University of the Republic in Uruguay (Protocol N∘ 070153000089-17). Adequate measures were taken to minimize pain, discomfort or stress to the animals. In addition, all efforts were made to use the minimum number of animals necessary to produce reliable scientific data.

### Surgical procedure

Following general anesthesia, the head was positioned in a stereotaxic frame and the skull was exposed. Stainless steel screw electrodes (1.4 mm diameter) were placed on the surface (above the dura matter) of different cortical areas including prefrontal, primary motor, primary somatosensory and posterior parietal cortices. Note that because the animals were not prepared specifically for this work, we did not analyze the same cortices in all of them. The electrodes were connected to a Winchester plug, which together with two plastic tubes were bonded to the skull with acrylic cement in order to maintain the animals’ head in fixed position without pain or pressure. After recovery from surgical procedures, they were adapted to the recording environment for a period of at least 2 weeks.

### Data acquisition and preprocessing

Experimental sessions of 4 h were conducted between 11 a.m. and 3 p.m. in a temperature-controlled environment (21–23 ∘C). During these sessions (as well as during the adaptation sessions), the animals’ head was held in a stereotaxic position by four steel bars that were placed into the chronically implanted plastic tubes, while the body rested in a sleeping bag (semi-restricted condition).

The ECoG activity was recorded with a monopolar (referential) configuration, utilizing a common reference electrode located in the left frontal sinus. The experiments on sleep and ketamine were performed on the same cats but not the same cortices were recorded as they were originally designed for different studies. The electromyogram (EMG) of the nuchal muscles, which was recorded by means of an acutely placed bipolar electrode, was also monitored. The electrocardiogram (ECG), by electrodes acutely placed on the skin over the pre-cordial region, and respiratory activity by means of a micro-effort piezo crystal infant sensor were also recorded. Each cat was recorded daily for ~30 days in order to obtain complete basal and treatment data sets. The animal retained muscular tone but hyperlocomotion was not observed in our experiments (Castro et al., 2019), nor in previous studies in cats (Issabeagloo et al., 2011), an increase in motor activity was also absent in semi-restricted condition, and ~5 min following the injection of ketamine the animals lay down on the floor unable to stand up (i.e., an ataxia-like effect), but responded to sound stimulus directing the gaze toward the sound source. In the absence of stimuli, the cats moved their head from one side to the other (i.e., a head-weaving-like behavior, described in rodents, and defined as stereotypies characterized as lateral side-to-side movement of the head without locomotion).

Bioelectric signals were amplified (×1,000), filtered (0.1 - 500 Hz), sampled (1,024 Hz, 2^16^ bits) and stored in a PC using the Spike 2 software (Cambridge Electronic Design).

Data were obtained after ketamine administration as well as during spontaneously occurring quiet W, LS, NREM sleep and REM sleep (Fig. 1). Five, 10, and 15 mg/kg i.m. of ketamine (Ketonal ®, Richmond Veterinaria S.A.) were administered to five animals in 4 different sessions. These three doses were administered in each animal in different experimental sessions performed in different days in a counterbalanced order. The scheme illustrated in figure 1B corresponds to one session, in which only one bolus of ketamine was administered. In each session, the animal was recorded in resting conditions for around 30 minutes and then the bolus of ketamine was injected.

After that, the recording continued for 4 hours. Ten minutes after the injection the cat received auditory stimulation (in 3 of the 5 cats). The different doses of ketamine were administered in different days leaving 3 or 4 days in between. Additionally, each different dose was repeated 4 times. In each session the whole experiment illustrated in figure 1B was repeated, therefore in total the experiments were repeated 12 times (4 per dose). Ketamine (50 mg/ml) was diluted in benzethonium chloride, hydrochloric acid, and water (solution for veterinary use). Basal recordings (without injections) were used as control. Sound stimuli were introduced ~30 min after the beginning of the recording sessions in drug-free condition, and 10 min after ketamine injection. These sound stimuli had the same characteristics as those used to induce active W (Castro et al., 2013). Sound stimuli was presented for a period of 300 s, and consisted of 60-100 dB SPL clicks, with variable frequency of presentation (1–500 Hz), modified at random in order to avoid habituation (Castro et al., 2013; Torterolo et al., 2003). The mentioned frequency refers to the frequency of presentation of the clicks and not the sound frequency. There were no frequency steps, and the SPL had no steps. Sound stimuli during 300 s were also performed 10 min after ketamine injection in three cats. The stimuli were square pulses produced with an electric stimulator connected to a speaker which emit them as clicks.

For preprocessing, sleep stages were scored off-line by visual inspection of 5-s epochs in Spike2 software, where the ECoG and electromyogram (EMG) were displayed simultaneously. In order to analyze LZs during sleep, a total of 300 artifact-free seconds data were selected from each behavioral state. Additionally, to study LZs during the Ketamine effect 300 s duration segments, with and without stimulation, were selected before and after ketamine administration.

After scoring, for both experiments, the selected epochs were exported to matlab for further preprocessing. The Matlab toolbox eeglab was used to filter the data (0.5-200 Hz band-pass). Each epoch was visually inspected, and those with gross artifacts (e.g. movements) were removed from the analysis.

### Lempel-Ziv complexity

In this study we used Lempel-Ziv (LZ) complexity to compute the complexity of measured neural signals (Lempel & Ziv, 1976). In particular, we used the LZ78 algorithm (Ziv & Lempel, 1978), which corresponds to the standard word-dictionary implementation: given a binary string, the algorithm scans it sequentially looking for distinct structures or “patterns.” The more diverse the binary string, the more patterns are included in the dictionary (a sequence containing only zeros or only ones would lead to the minimal number of patterns being obtained). The total number of these patterns is a measure of signal diversity.

To compute LZ from our experimental data, the recording of each channel was split into segments of 5120 samples (5s sampled at 1024Hz). Then, to generate a discrete sequence from a real-valued signal X of length *T*, X is detrended and binarized with a threshold of 0, and the resulting binary sequence is fed to the LZ78 algorithm. Finally, the resulting dictionary length *L* is normalized as

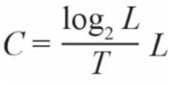

to yield a measure of complexity *C*.

Our choice on binarizing signals with a threshold of cero was driven by two factors: 1) LZ and related methods tend to be remarkably robust to the choice of discretisation procedure and number of bins (see e.g. discussion in Mediano (2020)); and 2) the Hilbert transform of a very broadband signal isn’t easily interpretable, and to create a meaningful analytic signal it would be necessary to bandpass-filter the data in a particular frequency band of interest. Given these two arguments, we reasoned that the added analysis complexity introduced by the filter parameters, frequency bands, etc, would not lead to substantially richer or more accurate results, and thus opted for the simple (yet probably effective) methodology.

### Statistics

One way ANOVA, with Tukey post-hoc test were used to compare LZ between sleep stages per cortex per animal (Fig. 2B) where Cohen’s d was used to address the size of the effect. Additionally, a multilevel approach as well as Bayesian Informational Criterion (BIC) were used to find the most likely explanatory model within the hierarchical model in the group statistical analysis comparing linear, quadratic and cubic models. For sleep study, the state of sleep was used as a fixed effect and the cat identity as a random effect. The same type of approach was used to study the ketamine effect among different cortices under control and stimulus conditions. In this case the dose and stimulus (if present) were used as fixed effects; and cat identity and session as random effects. The interaction between stimuli and ketamine dose was also included in the model when studying the modulation by stimulus. All models were estimated via restricted maximum likelihood, using the open-source packages lme4 v.1.1-21 (Bates et al., 2015) and lmerTest v.3.1-1 (Kuznetsova et al., 2017) on R v.3.6.1.

## Data and materials availability

The code for computation of LZ used for analysis is available in GitHub at the following link: https://gitlab.com/CPasco83/sleep-and-ketamine. Data is available upon reasonable request to the authors.

## Authors’ Contribution

P.T. and T.B. designed the study. C.Z.S. and P.T. performed the experiments and collected the data. C.P. and P.M. analyzed data; M.P. and P.M. wrote LZ codes; C.P. wrote the manuscript; all authors participated in the interpretation of results and revision of the manuscript, and approved the final version of the manuscript. P.T., T.B. and D.B. provided the financial support.

## Acknowledgements

This study was supported by the “Programa de Desarrollo de Ciencias Básicas, PEDECIBA” and the “Comisión Sectorial de Investigación Científica” (CSIC) I + D-2020-393 grant from Uruguay. PAM and DB are funded by the Wellcome Trust (grant no. 210920/Z/18/Z).

## Abbreviations

BIC: Bayesian information criterion
BF: Bayes Factors
DMT: N,N-Dimethyltryptamine
ECG: Electrocardiogram
ECoG: Electrocorticogram
EEG: Electroencephalography
EMG: Electromyogram
ERPs: Evoked related potentials
GABA: Gamma aminobutyric acid
HCN channels: Hyperpolarization-activated cyclic nucleotide-gated channels
LS: Light sleep
LSD: Lysergic acid diethylamide
LZ: Lempel Ziv complexity
NMDA: N-methyl-D-aspartate
NREM: Non-rapid eyes movement
PCI: Perturbational complexity index
PCI^ST^: PCI state-transition
REM: Rapid eye movements
SWS: Slow wave sleep
TMS: Transcranial magnetic stimulation
5-HT: 5-hydroxytryptamine

