## Supplementary Figure 1 for "Ketamine and sleep modulate neural complexity dynamics in cats"

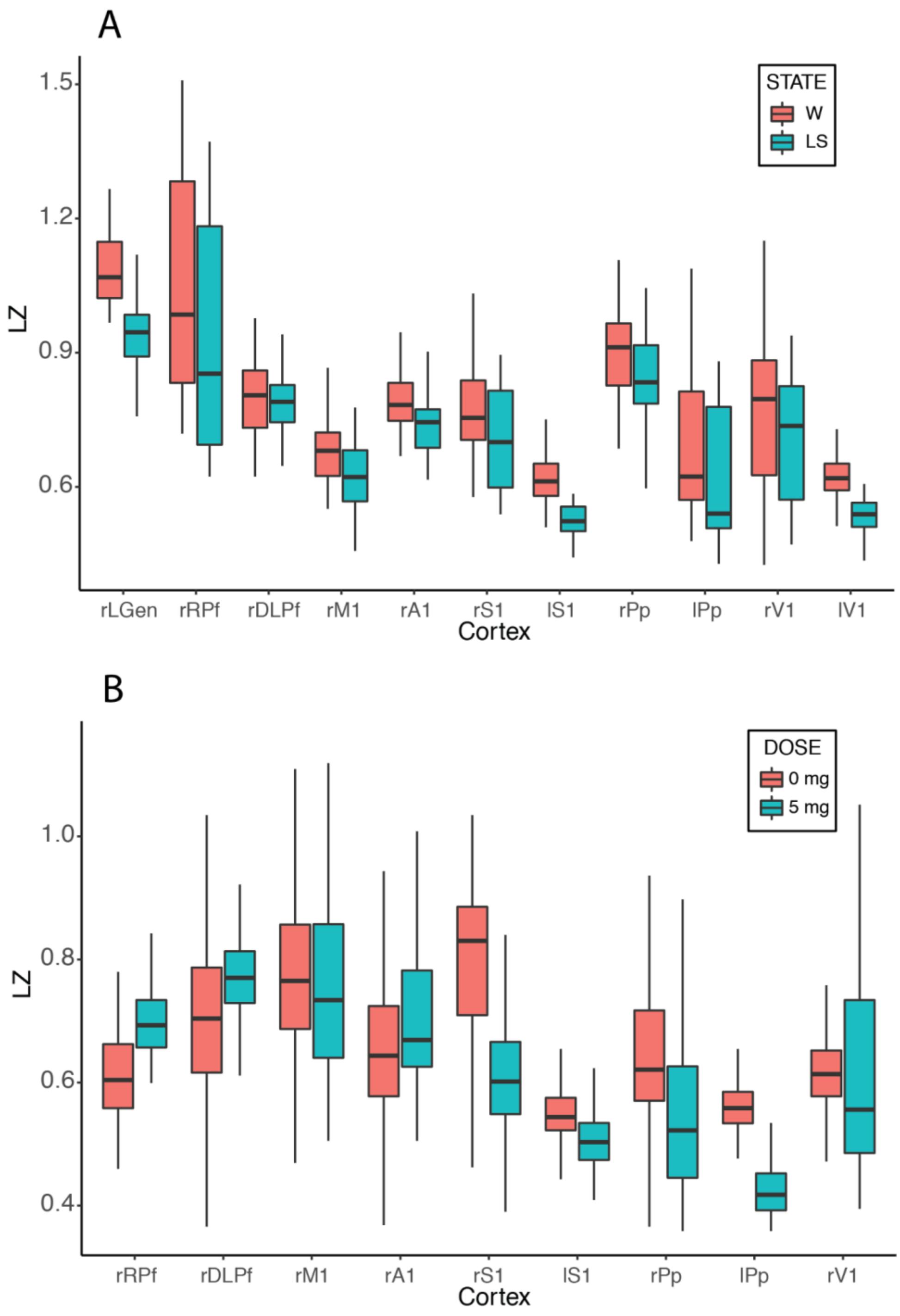


SUPPLEMENTARY FIGURE 1. Comparison between LZ in wakefulness and LS (A), and wakefulness without ketamine vs. 5 mg of Ketamine (B) for the different cortices. RPf, rostral prefrontal cortex; DLPf, dorsolateral prefrontal cortex; M, primary motor cortex; S, primary somatosensory cortex; A, primary auditory cortex, Pp, posterior parietal cortex; V, visual cortex. “l” indicates left side, “r” right side and “1” indicates “primary cortex”.
